# Endothelial cells secreted ET-1 augments DN via inducing EM accumulation of MCs in ETBR^−/−^ mice

**DOI:** 10.1101/303768

**Authors:** Hong-hong Zou, Li Wang, Yun-feng Shen, Xiao-xu Zheng, Gao-si Xu

## Abstract

ETBR deficiency may contribute to the progression of DN in a STZ model, but the underlying mechanism is not fully revealed. In this study, STZ-diabetic ETBR^−/−^ mice was characterized by increased serum creatinine, urinary albumin and ET-1 expression, and enhanced glomerulosclerosis compared with STZ-diabetic WT mice. HG conditioned media of ETBR^−/−^ endothelial cells promoted MC proliferation and upregulated ECM-related proteins, and ET-1 knockout in endothelial cells or inhibition of ET-1/ETAR in MC suppressed MC proliferation. ET-1 was over-expressed in ETBR^−/−^ endothelial cells and was regulated by NF-kapapB pathway. And ET-1/ETBR suppressed NF-kappaB via eNOS to modulate ET-1 in endothelial cells. Furthermore, ET-1/ETAR promoted RhoA/ROCK pathway in MC, and accelerated MC proliferation and ECM accumulation. *In vivo* experiments proved ETBR^−/−^ mice inhibited NF-kappaB pathway to ameliorate DN and eNOS mice had similar results. Hence, in HG-exposed ETBR^−/−^ endothelial cells, suppression of ET-1/ETBR activated NF-kappaB pathway via inhibiting eNOS to secrete large amount of ET-1. Due to the communication between endothelial cells and MCs, ET-1/ETAR in MC promoted RhoA/ROCK pathway to accelerate MC proliferation and ECM accumulation.

## Introduction

Diabetic nephropathy (DN), characterized by renal inflammation, urinary albumin, decreased glomerular filtration rate, glomerulosclerosis and tubulointerstitial fibrosis, is a leading cause of end-stage renal disease (ESRD) [1, 2]. Thus, exploring the mechanisms that mediate DN is critical for the prevention and treatment strategies for alleviating DN. Mesangial cells (MC) are critical in maintaining mesangial matrix homeostasis, regulating glomerular filtration rate, and keeping normal glomerular function via producing cytokines, metalloproteinases and extracellular matrix (ECM) [3]. Researchers have found that MC proliferation and ECM accumulation were main contributing factors to glomerulosclerosis and tubulointerstitial fibrosis, which were important characters of DN [4, 5]. In high-glucose (HG) condition, fibronectin, collagen IV, and plasminogen activator inhibitor were observed to be upregulated in MCs [6]. Connective tissue growth factor (CTGF), a growth factor produced by activated MCs, played a key role in the pathogenesis of DN and was also upregulated in DN[7]. Hence, glomerulosclerosis is closely associated with dysfunction of MCs under HG condition.

Endothelin-1 (ET-1) is mainly expressed by endothelial cells and plays important role in DN [8, 9]. ET-1 exerts its effects by binding to one of two endothelin receptor subtypes, namely Endothelin A Receptor (ETAR) and Endothelin B Receptor (ETBR). Researchers have found ETBR was mainly distributed in vascular and glomerular endothelial cells, whereas ETAR was mainly distributed in smooth muscle cells and MCs [10]. Reports have shown that MCs and endothelial cells interacted in the kidney [11, 12], however, studies mainly focused on the ability of MCs to regulate the synthesis of ET-1 by endothelial cells, the effect of endothelial cells on MCs is rarely discussed. Based on ET-1 mainly expressed by endothelial cells and the crosstalk between endothelial cells and MC, we speculated that ETBR knockout in endothelial cells might have vital function in MC proliferation and ECM accumulation.

Given the previous report of diabetic ETBR-deficient (ETBR^−/−^) rats caused progressive renal failure [13], we established streptozotocin (STZ)-diabetic ETBR^−/−^ mice model to explore the exact mechanism of ETBR^−/−^ mice in the acceleration of DN. Our results showed that STZ-diabetic ETBR^−/−^ mice enhanced glomerulosclerosis and had higher levels of renal damage signs (serum creatinine and urinary albumin) and increased mRNA and protein levels of ET-1 *in vivo*. Under high glucose condition, ETBR^−/−^ endothelial cells secreted large amount of ET-1, thus to promote ECM accumulation of MC *in vitro*.

## Materials and methods

### Animals and establishment of diabetic mice

Male C57BL/6 (wild-type, WT) mice and ETBR^−/−^ mice (aged 7 weeks) were intraperitoneally injected with 50 mg/kg STZ everyday for five days to establish STZ-diabetic mice model. Diabetic mice were confirmed two weeks after the initial intraperitoneal injection with the criteria of a blood glucose >16 mmol/L. WT mice and ETBR^−/−^ mice (n=5/group) were used as control and received an intraperitoneal injection of 0.1 M citrate buffer (pH 4.5) everyday. All mice were kept in metabolic cages and had a standard diet (0.2% sodium) with no limitation to water. Before killing, blood was collected from retro-orbital vein plexus, after 4 °C for the night and centrifugation for 10 min at 3000×g, serum was obtained and stored at −20. Urine was also collected from the mice. Ten weeks later, all the mice were sacrificed to detect the indices. kidneys were collected for further use and weighed. The animal study was approved by the Ethic Committee of the Second Affiliated Hospital of Nanchang University.

*In vivo* experiment, C57BL/6 mice, endothelial nitric oxide synthase^−/−^ (eNOS^−/−^) mice, and ETBR^−/−^ mice (n=3/group) were intraperitoneally injected with 50 mg/kg STZ everyday for five days to establish STZ-diabetic mice model. Diabetic mice were confirmed two weeks after the initial intraperitoneal injection with the criteria of a blood glucose >16 mmol/L. Bay 11-7082 (1mg/kg) was injected into mice between seventh and 10th weeks after STZ treatment.

### Serum and urine detection

The levels of serum glucose, serum creatinine, urinary albumin were detected by 7600 automatic biochemical analyzer (HITACHI, Japan). ET-1 concentration was measured by ET-1 ELISA kit (Shanghai Jingkang Biotechnology, China) according to the manufacturer’s instructions.

### Histology

After fixation of the kidney, the slices were embedded in paraffin. Sections of 3μm were stained by periodic acid-Schiff (PAS) and hematoxylin. Glomerulosclerosis was defined by the presence of PAS-positive material within the glomeruli. Twenty glomeruli specimens in each group were used to observe the glomerulosclerosis. The scoring guidelines of the proportion of PAS-positive material within each glomerulus are: 1, a proportion <25%; 2, a proportion of 25%–50%; 3, a proportion of 50%–75%; 4, a proportion > 75%. The average score assigned to all glomeruli was defined as glomerulosclerosis index.

### Cell culture and transfection

Primary endothelial cells were obtained from the following procedures: pronase (Roche, Switzerland) followed by collagenase (Roche, Switzerland) were used for in situ perfusion of the liver from ETBR^−/−^ mice. Discontinuous density gradient of Accudenz (Accurate Chemical and Scientific, Canada) was used to layer cell suspensions. Endothelial cells were present in the lower layer and further purified by centrifugal elutriation (18 ml/min flow). Cells were cultured in DMEM/F-12 (Gibco, USA) containing 20% fetal bovine serum (Gibco, USA). Endothelial cells after the third passage were probed with anti-CD31 (BD, USA) to confirm the purity of primary endothelial cells was greater than 95%.

Mouse glomerular mesangial cell (MC) line SV40 MSE13 was purchased from Cell bank of Chinese Academy of Sciences (Shanghai, China) and cultured in DMEM/F-12 (Gibco, USA) containing 10% FBS (Gibco, USA), 14mM HEPES (Gibco, USA), 150mg/L L-glutamine (Sinopharm Chemical Reagent, China) and 1.5g/L NaHCO_3_ (Sinopharm Chemical Reagent, China) in 5% CO_2_ incubator under 37°C.

Cells were seeded into a culture plate and grown to 80% confluence for cell transfection. si-ET-1, si-p65, si-ETBR, and si-eNOS were synthesised by GENECHEM (Shanghai, China), and were transfected into cells using Lipofectamine 2000 (Invitrogen, USA).

### Western Blot

Proteins from kidney, endothelial cells or MC were isolated from using RIPA buffer (Thermo Scientific, USA). 50 μg of protein samples were isolated in 12% SDS-polyacrylamide gel electrophoresis (SDS-PAGE), and transferred to polyvinylidene difluoride (PVDF) membranes (Invitrogen, USA) by electroblotting. The membranes were blocked in 5% non-fat dried milk for 60 min at room temperature. For the detection of RhoA, membrane and cytosolic proteins were separated on SDS-PAGE. The membranes were probed with first primary antibody anti-ETBR (Abcam, USA), anti-ETAR (Abcam, USA), anti-p-eNOS (Millipore, USA), anti-eNOS (Invitrogen, USA), anti-p-p65 (Invitrogen, USA), anti-p65 (Invitrogen, USA), anti-CTGF (Abcam, USA), anti-RhoA (Abcam, USA), anti-collagen IV (Abcam, USA), anti-Fibronectin (Abcam, USA), anti-p21 (Abcam, USA) and anti-β-actin (Invitrogen, USA) and incubated at 4°C overnight. After washing with PBST, membrane was cultivated with secondary antibody for 60 min at room temperature. β-actin was used as internal control.

### Quantitative real-time PCR (qRT-PCR)

To quantify mRNA expression of ET-1, eNOS, ETAR and ETBR, we conducted qRT-PCR. Total RNA from kidney tissue, endothelial cells or MC were isolated by TRIzol Reagent (Invitrogen, USA) according to manufacturer’s instructions, and was transcribed to cDNA with iScript cDNA Synthesis kit (Bio-Rad, USA). PCR was performed as follows: 94°C for 5 min, then 35 cycles of denaturation at 94°C for 30 s, annealing at 64°C for 30 s and extension at 72°C for 120 s at QuantStudio® 3 RCR Real-Time PCR systems (Applied Biosystems, USA). The relative ET-1, eNOS, ETAR and ETBR expressions were determined by via the comparative 2-^ΔΔCq^ method.

### MTT assay

3-(4,5-dimethylthiazol-2-yl)-2,5-diphenyltetrazolium bromide (MTT) assay was used to detect the proliferation of MC. 1× 10^5^ MC were seeded into a 96-well plate. After overnight incubation with different treatment, 20 μl MTT (5 mg/mL; Invitrogen, USA) was added to each well and cultured for 2 h. Then, cells were lysed using dimethylsulfoxide (150 μl/well; Sinopharm Chemical Reagent, China). The optical density was read at 570 nm.

### Apoptosis assay

After washing with cold phosphate-buffered saline (PBS), 5 μl of annexin-V-FITC (Beyotime Biotechnology, China) was added to the MCs and incubated at room temperature for 15 min. Then, 10 μl propidium iodide (PI) was added before flow cytometry analysis. Apoptosis were measured using a FACS Calibur flow cytometer (BD, USA) and analyzed by Cell Quest pro software.

### Cell cycle analysis

After 72 h of treatment, MCs were harvested by trypsinization and washed with cold PBS for two times. Then, MCs were fixed with 75% alcohol for 12 h at 4°C. After washing with cold PBS, cells were treated with 50 ug/mL RNase for 30 min at 37°C, then stained with 50 μg/mL PI for 30 min at 4°C in the dark before being analyzed using a FACS Calibur flow cytometer (BD, USA). Cells (2×10^6^) were detected for each sample. Cell cycle was analyzed by FlowJo software.

### Reporter gene assay

Endothelial cells were transfected with a reporter plasmid using Lipofectamine LTX Reagent (Invitrogen, USA) according to the manufacture’s instruction, and firefly luciferase reporter was used to measure ET-1 promoter activity. After 48 h of transfection, dual-Luciferase Reporter Assay System (Promega, USA) was used to measure reporter activities.

### Statistical analysis

SPSS software (version 18.0) was used for data analysis, and the result was expressed as mean ± standard deviation (SD). One-way ANOVA and t test were used for the data analysis, with p < 0.05 considered statistically significant.

## Results

### Severer diabetic nephropathy in ETBR^−/−^ mice

We observed that STZ-diabetic mice had reduced body weight, increased kidney weight and increased kidney/body weight ratio (Figure 1A, 1B and 1C). And there were no significant differences in body weight and kidney weight between STZ-diabetic ETBR^−/−^ mice and STZ-diabetic WT mice. As shown in Figure 1D, STZ-diabetic mice had higher serum glucose level than control mice. In addition, STZ-diabetic mice had higher serum creatinine and urinary albumin level than control mice. And serum creatinine and urinary albumin level in STZ-diabetic ETBR^−/−^ mice were significantly higher than that of STZ-diabetic WT mice (Figure 1E and 1F). Histopathological analysis showed that enlargement of glomeruli was observed in STZ-diabetic mice, and enhanced glomerulosclerosis was present in STZ-diabetic ETBR^−/−^ mice (Figure 1G).

**Figure 1.**
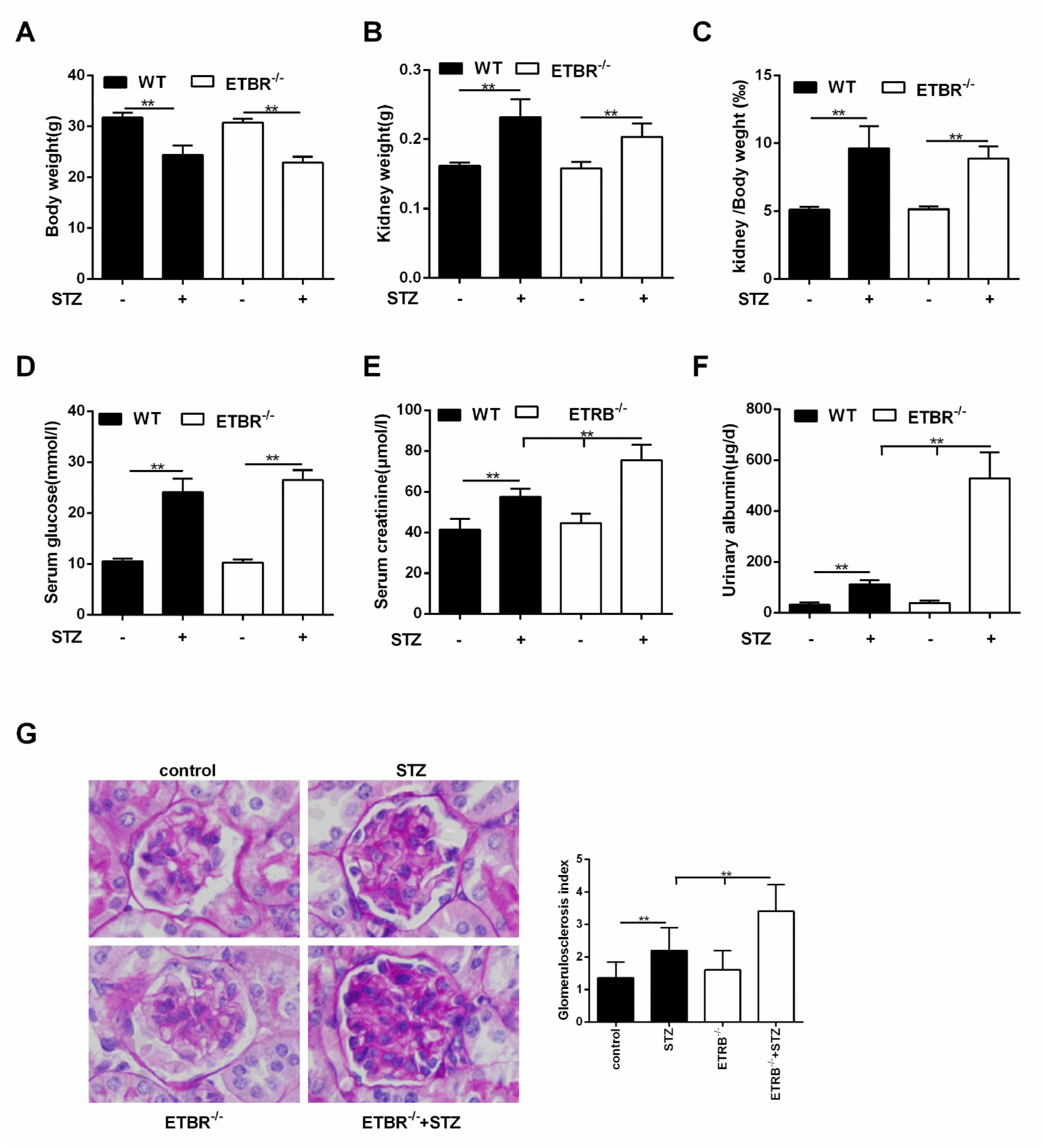
Severer diabetic nephropathy in ETBR^−/−^ mice. A-C. Compared with control mice, STZ-diabetic mice had reduced body weight, increased kidney weight and increased kidney/body weight ratio. And there were no significant differences in body weight and kidney weight between STZ-diabetic ETBR^−/−^ mice and STZ-diabetic WT mice. D. STZ-diabetic mice had higher serum glucose level than control mice. E-F. STZ-diabetic mice had higher serum creatinine and urinary albumin level than control mice. And serum creatinine and urinary albumin level in STZ-diabetic ETBR^−/−^ mice were significantly higher than that of STZ-diabetic WT mice. G. Periodic acid-Schiff (PAS) and hematoxylin staining showed that enlargement of glomeruli was observed in STZ-diabetic mice, and enhanced glomerulosclerosis was present in STZ-diabetic ETBR^−/−^ mice. **p<0.01 versus control mice.

### ET-1 expression was increased in STZ-diabetic ETBR^−/−^ mice

As shown in Figure 2A, protein levels of CTGF, ETAR and p-p65 in STZ-diabetic ETBR^−/−^ mice was higher than STZ-diabetic WT mice, whereas phosphorylation-eNOS (p-eNOS) protein level was lower than STZ-diabetic WT mice. Besides, we found mRNA (Figure 2B) and protein (Figure 2C and 2D) expressions of ET-1 in STZ-diabetic ETBR^−/−^ mice were higher than STZ-diabetic WT mice.

**Figure 2.**
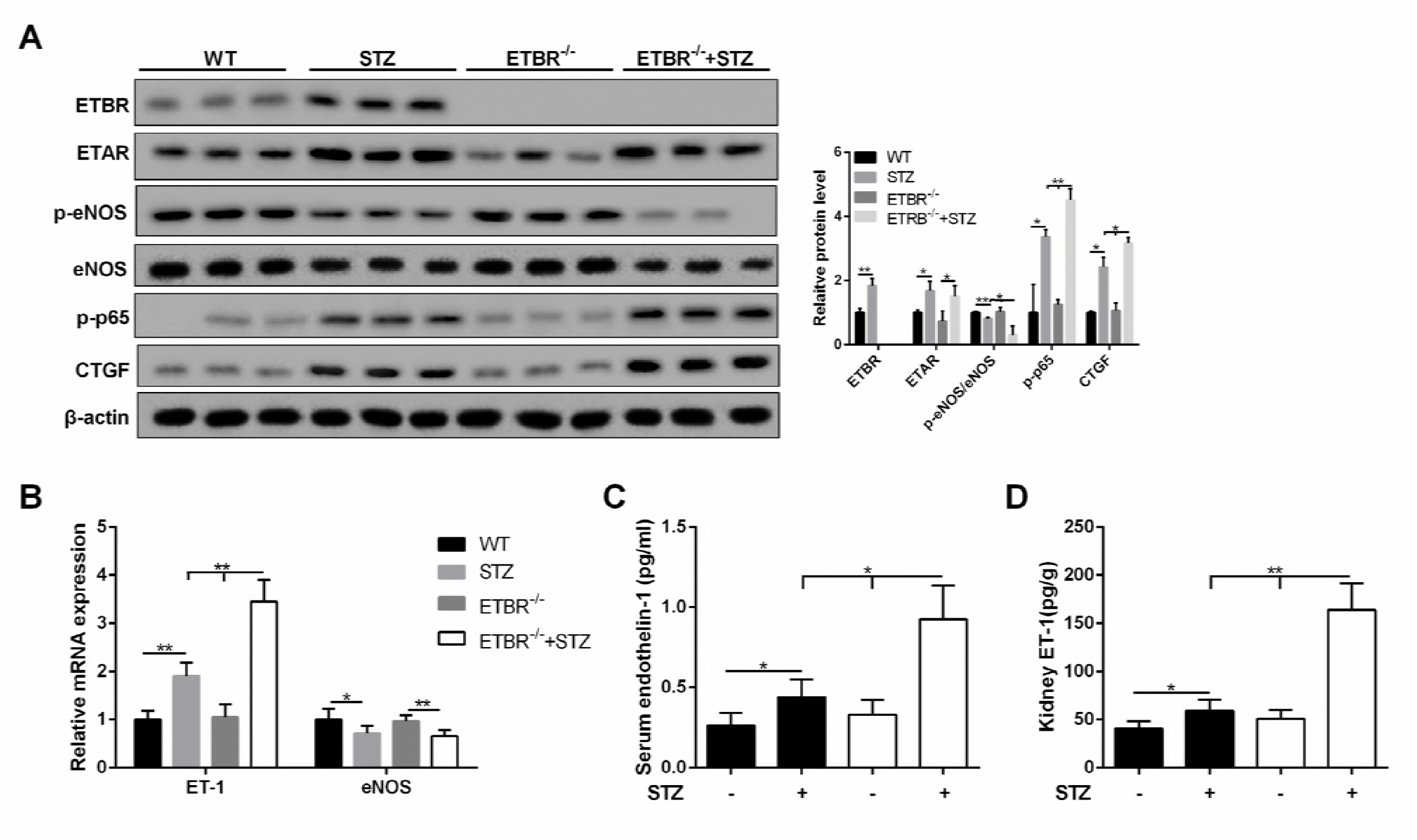
ET-1 expression in STZ-diabetic ETBR^−/−^ mice was higher than STZ-diabetic WT mice. A. CTGF protein level in STZ-diabetic ETBR^−/−^ mice was higher than STZ-diabetic WT mice, whereas p-eNOS protein level was lower. *p<0.05 versus WT mice or STZ-diabetic mice. **p<0.01 versus WT mice or STZ-diabetic mice. B-D. mRNA and protein expressions of ET-1 in STZ-diabetic ETBR^−/−^ mice were higher than STZ-diabetic WT mice. *p<0.05 versus WT mice or STZ-diabetic mice or STZ-diabetic WT mice. **p<0.01 versus WT mice or STZ-diabetic mice.

### High-glucose conditioned media (CM) of ETBR^−/−^ endothelial cells promoted MC proliferation and ECM formation

After 24 h of high-glucose treatment, protein level of ET-1 in primary endothelial cells of ETBR^−/−^ mice was significantly higher than that of WT mice (Figure 3A). In this study, CM of ETBR^−/−^ endothelial cells under normal or HG condition was used to cultivate MCs. HG-treated CM of ETBR^−/−^ endothelial cells promoted proliferation of MC (Figure 3B). HG-treated CM increased RhoA level on MC membrane and facilitated Collagen IV secretion, and the strengthen effect of ETBR^−/−^ CM on MC was stronger than WT CM (Figure 3C). Later, we explored why HG-treated CM promoted MC proliferation and ECM formation. After ET-1 knockout in endothelial cells, HG-treated CM significantly inhibited MC proliferation and Collagen IV formation, and the result was similar in ABT-627 group (inhibition of ET-1/ETAR pathway in MC) (Figure 3D and 3E). There was no significant difference in the result between A192621 group (inhibition of ET-1/ETBR pathway in MC) and ET-1 normal group. These results pointed out whether glomerulosclerosis in diabetic mice was related with ET-1 secretion, whether increased glomerulosclerosis index in ETBR^−/−^ mice was associated with higher ET-1 secretion in ETBR^−/−^ endothelial cells, and why there was higher ET-1 secretion in ETBR^−/−^ endothelial cells. These questions would be explored in the following experiments.

**Figure 3.**
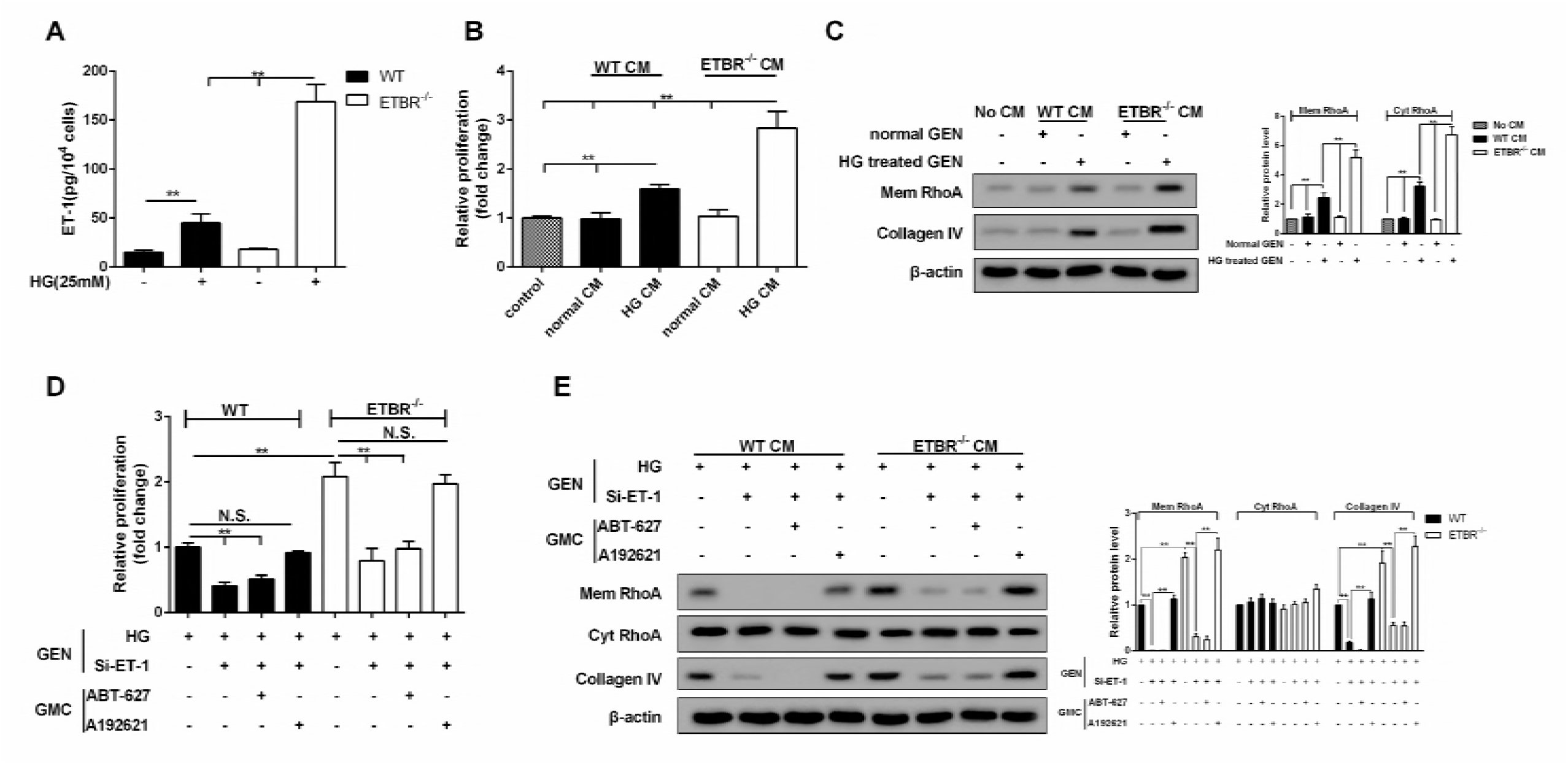
High-glucose conditioned media (CM) of ETBR^−/−^ endothelial cells promoted mesangial cell (MC) proliferation and ECM formation. A. After 24 h of high-glucose (25mM) treatment, ET-1 level in primary endothelial cells of ETBR^−/−^ mice was significantly higher than that of WT mice. **p<0.01 versus control or HG WT. B-C. CM of ETBR^−/−^ endothelial cells was used to cultivate MC. MC in control group was cultured in normal medium. HG-treated CM of ETBR^−/−^ endothelial cells promoted proliferation of MC. HG-treated CM increased RhoA level on MC membrane and facilitated Collagen IV secretion, and the strengthen effect of ETBR^−/−^ CM was stronger than WT CM. **p<0.01 versus control or normal WT CM or HG WT CM or normal ETBR^−/−^ CM. D-E. The reason of endothelial cell CM on the promotion of MC proliferation and ECM formation was explored. After ET-1 knockout in endothelial cells, HG-treated CM significantly inhibited MC proliferation and Collagen IV formation, and the result was similar in ABT-627 group (inhibition of ET-1/ETAR pathway in MC). There was no significant difference in the result between A192621 group (inhibition of ET-1/ETBR pathway in MC) and ET-1 normal group. **p<0.01 versus si-ET-1 or si-ET-1+ABT-627 or si-ET-1+A192621.

### ET-1 modulated MC proliferation and ECM through RhoA/Rho-kinase (ROCK) pathway

As shown in Figure 4A, MC proliferation was promoted with the increase of ET-1 concentration under HG serum-free condition. Under HG serum-free condition, ECM-related proteins (Collagen IV, Fibronectin and CTGF) and RhoA on MC membrane were also up-regulated with the increase of ET-1 concentration (Figure 4B). In addition, Y-27632 (RhoA/ROCK inhibitor) suppressed MC proliferation and promoted cell apoptosis, and cell cycle was arrested at G0/G1 phase with the treatment of ET-1 (Figure 4C, 4D and 4E). Moreover, Y-27632 increased p21 level (a regulator of cell cycle progression at G1 phase). Therefore, MC cycle was arrested, the proliferation was decreased, and ECM-related proteins were down-regulated with the treatment of ET-1 (Figure 4F).

**Figure 4.**
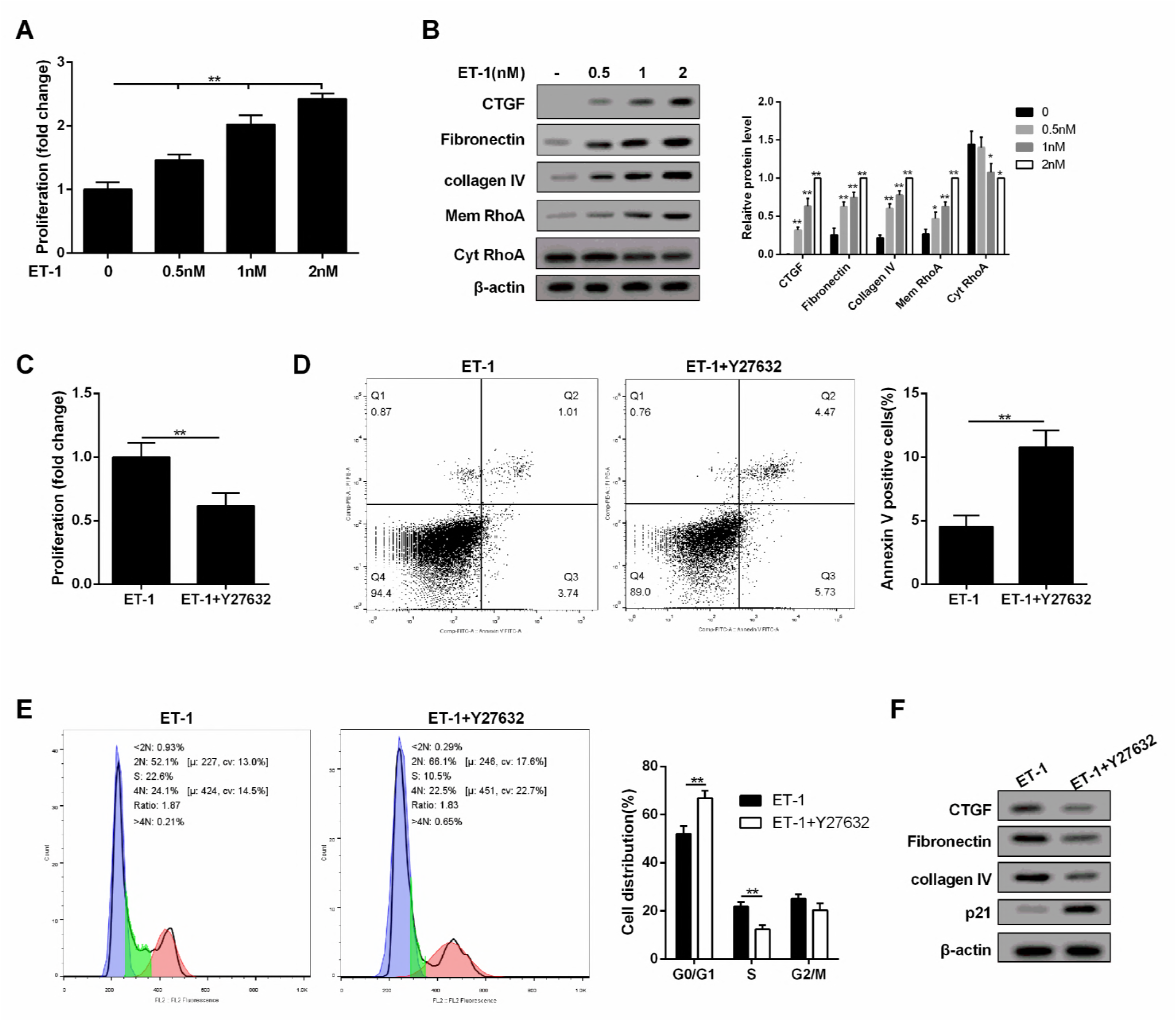
ET-1 modulated MC proliferation and ECM through RhoA/ROCK pathway. A. Under HG serum-free condition, MC proliferation was promoted with the increase of ET-1 concentration. **p<0.01 versus 0nM. B. Under HG serum-free condition, ECM-related proteins (Collagen IV, Fibronectin and CTGF) and RhoA on MC membrane were upregulated with the increase of ET-1 concentration. ET-1 (1nM, 2500pg/ml) was used for the following experiments. **p<0.01 versus 0nM. C-E. Under HG serum-free condition+ET-1 treatment, Y-27632 (RhoA/ROCK inhibitor) suppressed cell proliferation and promoted cell apoptosis, and cell cycle was arrested at G0/G1 phase. **p<0.01 versus ET-1. F. Under HG serum-free condition+ET-1 treatment, Y-27632 (RhoA/ROCK inhibitor) increased p21 level, therefore, MC cycle was arrested, the proliferation was decreased, and ECM-related proteins were downregulated.

### ET-1 promoted RhoA/ROCK pathway in MC through ETAR

To explore whether ET-1 promoted RhoA/ROCK pathway through ETAR or ETBR, we administered ET-1/ETAR pathway inhibitor and ET-1/ETBR pathway inhibitor (ABT-627 and A192621) to MCs. We found inhibition of ETAR pathway inhibited MC proliferation, RhoA/ROCK and ECM-related proteins, and increased cell apoptosis, while inhibition of ETBR pathway didn’t affect cell growth, RhoA/ROCK and ECM-related proteins (Figure 5A, 5B and 5C). Besides, expression quantity from gene transcription level of ETAR in MC was higher than ETBR (Figure 5D). In ET-1 treated MC, ETAR expression on MC membrane was increased, while ETBR expression didn’t change, which increased the probability of combining ET-1 with ETAR. Hence, ET-1/ETBR pathway was not the dominant in MC. Under the stimulation of ET-1, the combination of ET-1 with ETAR activated the RhoA/ROCK pathway, thus affecting the growth of MC and the secretion of ECM. Therefore, we speculated although ETBR was knocked out in ETBR^−/−^ mice, the effect of ET-1 stimulation on ETBR^−/−^ MC was similar as that on WT MC.

**Figure 5.**
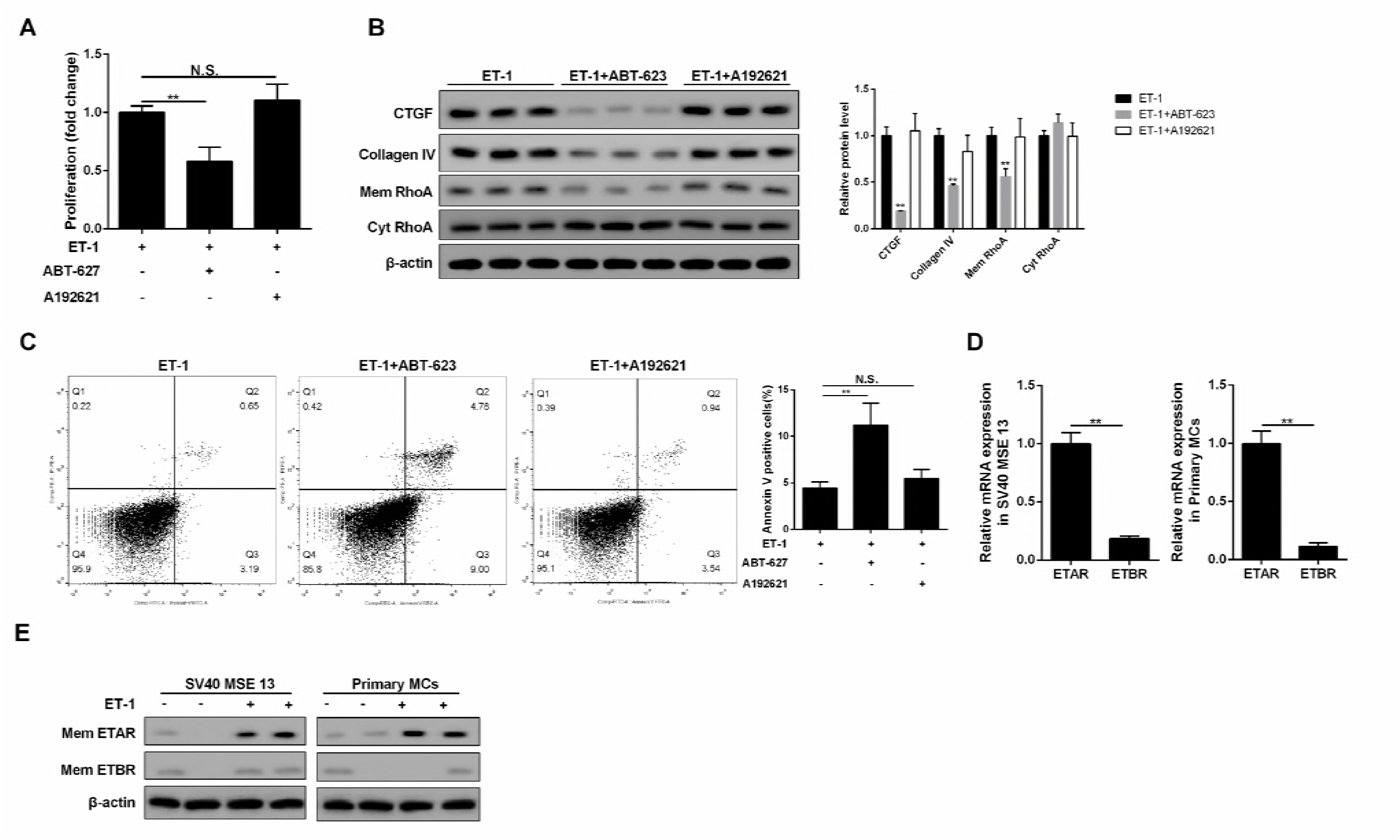
ET-1 promoted RhoA/ROCK pathway in MC through ETAR. A-C. Inhibition of ETAR pathway inhibited cell proliferation, RhoA/ROCK and ECM-related proteins, and increased cell apoptosis, while inhibition of ETBR pathway didn’t affect cell growth, RhoA/ROCK and ECM-related proteins. **p<0.01 versus ET-1. D. Expression quantity from gene transcription level of ETAR in MC was higher than ETBR. **p<0.01 versus ETAR. E. In ET-1 treated MC, ETAR expression on MC membrane was increased, while ETBR didn’t change, which increased the probability of combining ET-1 with ETAR.

### ET-1 was over-expressed in ETBR^−/−^ endothelial cells and was regulated by NF-kapapB pathway

As shown in Figure 6A, ET-1 expression was increased with time in WT and ETBR^−/−^ endothelial cells under the HG condition, and there was significant difference in ET-1 expression between ETBR^−/−^ and WT endothelial cells until 20 h, which indicated that endothelial cells grew fast between 16 h and 24 h. Besides, there was significant difference in ET-1 production rate between ETBR knockout and WT endothelial cells since 16 h, and the difference increased with time (Figure 6B). We found mRNA expression of ET-1 in WT endothelial cells reached the highest at 16 h (Figure 6C), and mRNA expression of ET-1 was increased in ETBR^−/−^ endothelial cells within 24 h (Figure 6D). Protein level of p-p65 in WT endothelial cells reached the highest at 12 h, and protein level of p-p65 was increased in ETBR^−/−^ endothelial cells within 24 h (Figure 6E). Therefore, we speculated whether Nuclear factor-kappaB (NF-kappaB) pathway was related with ET-1 mRNA level. As shown in Figure 6F and 6G, NF-kappaB inhibitor (Bay) decreased mRNA expression of ET-1 in WT or ETBR^−/−^ endothelial cells, indicating ET-1 mRNA was regulated by NF-kappaB. We also found extracellular secretion of ET-1 was regulated by NF-kappaB (Figure 6H). Importantly, Luciferase assay showed that Bay and si-p65 significantly decreased ET-1 promoter activity, which indicated that NF-kappaB regulated promoter of ET-1 at transcription level (Figure 6I). These findings indicated that NF-kappaB regulated promoter of ET-1 at transcriptional level, thus regulating ET-1 expression at mRNA and protein levels.

**Figure 6.**
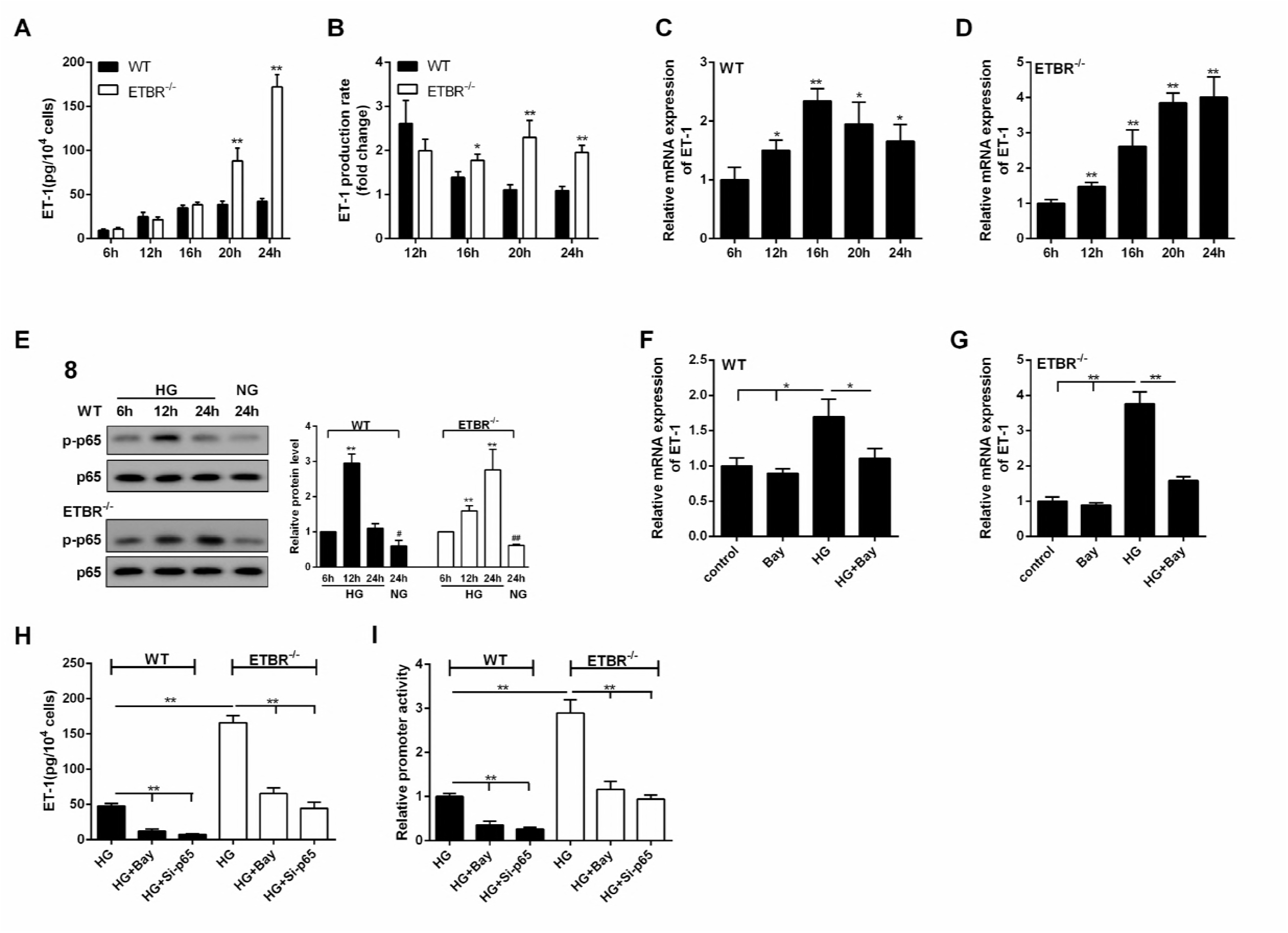
ET-1 was overexpressed in ETBR knockout endothelial cells and was regulated by NF-kapapB pathway. A. Under the HG condition, ET-1 expression was increased with time in WT and ETBR knockout endothelial cells, and there was significant difference in ET-1 expression between ETBR knockout and WT endothelial cells until 20 h, which indicated that endothelial cells grew fast between 16 h and 24 h. **p<0.01 versus WT. B. There was significant difference in ET-1 production rate between ETBR knockout and WT endothelial cells since 16 h, and the difference increased with time. B.ET-1 production rate (n) = ET-1(n)/ET-1(n-4). *p<0.05 versus WT. **p<0.01 versus WT. C. mRNA expression of ET-1 in WT endothelial cells reached the highest at 16 h. *p<0.05 versus 6 h. **p<0.01 versus 6h. D. mRNA expression of ET-1 was increased in ETBR knockout endothelial cells within 24 h. **p<0.01 versus 6h. E. Protein level of p-p65 in WT endothelial cells reached the highest at 12 h. And protein level of p-p65 was increased in ETBR knockout endothelial cells within 24 h. F-G. In WT or ETBR knockout endothelial cells, NF-kappaB inhibitor (Bay) decreased mRNA expression of ET-1, indicating ET-1 mRNA was regulated by NF-kappaB. H. Extracellular secretion of ET-1 was regulated by NF-kappaB. I. Luciferase assay showed that NF-kappaB regulated promoter of ET-1 at transcription level.

### ET-1/ETBR suppressed NF-kappaB via eNOS to modulate ET-1

In order to explore the mechanism of ET-1 in endothelial cells, A192621 and si-ETBR were used to inhibit ET-1/ETBR pathway. Under the HG condition, inhibition of ETBR pathway increased ET-1 secretion and mRNA expression in WT endothelial cells, and the effect of si-ETBR knockout was similar as inhibition of ETBR pathway (Figure 7A and 7B). There was no significant difference in ET-1 secretion between ETAR inhibition group and control group. We observed A192621 or si-ETBR suppressed protein level of p-eNOS and increased protein level of p-p65 (Figure 7C). Therefore, eNOS might be related with p-p65 expression. We further silenced eNOS, and found si-eNOS significantly increased protein level of p-p65, indicating p-p65 was negatively regulated by eNOS (Figure 7D). After further treatment of Bay, mRNA expression and secretion of ET-1 were significantly suppressed (Figure 7E and 7F). Hence, we proved ET-1/ETBR suppressed NF-kappaB via eNOS to modulate ET-1 expression.

**Figure 7.**
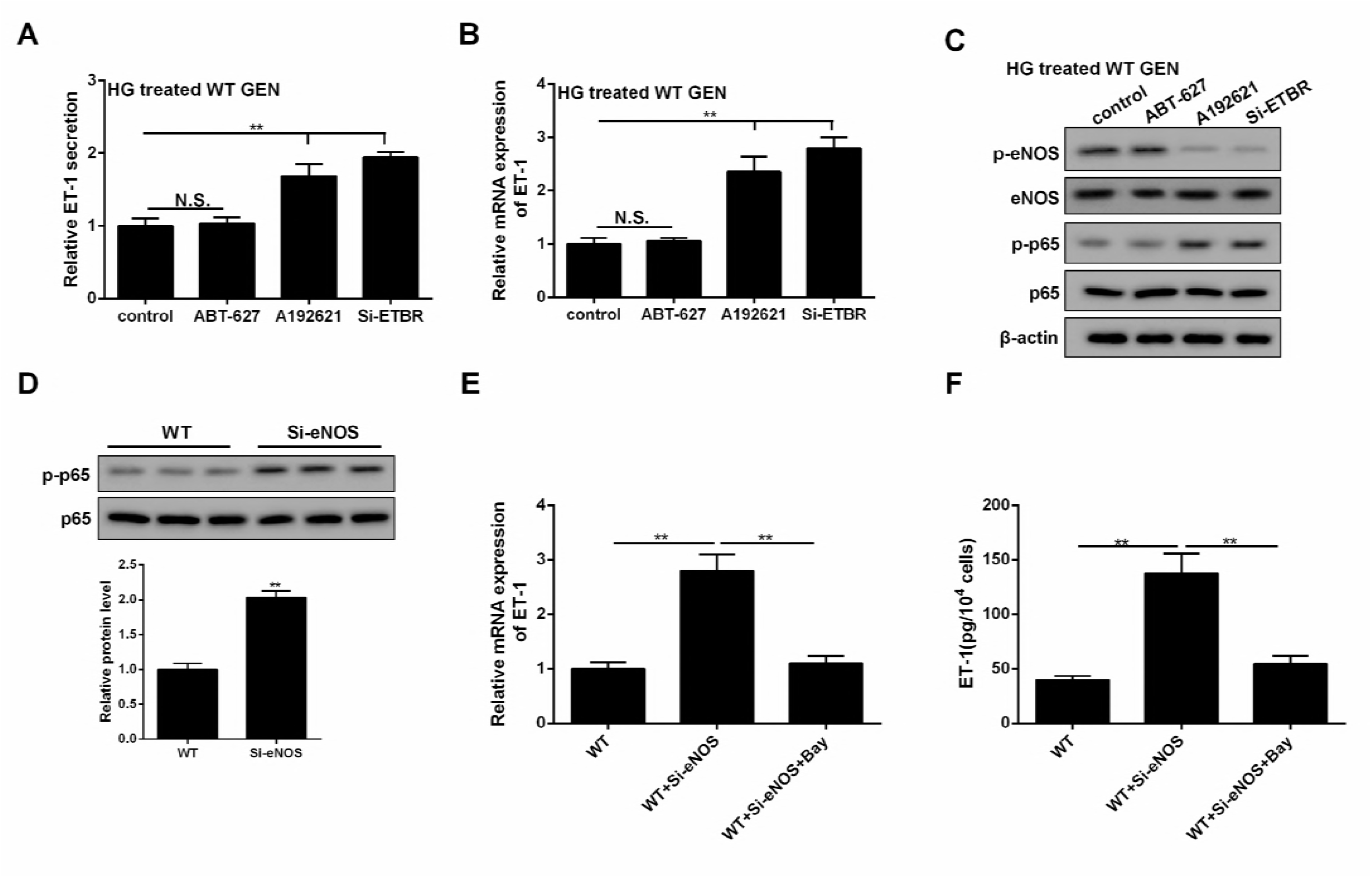
ET-1/ETBR suppressed NF-kappaB via eNOS to modulate ET-1. A-B. Under the HG condition, inhibition of ETBR pathway increased ET-1 secretion and mRNA expression in WT endothelial cells, and the effect of si-ETBR knockout was similar as inhibition of ETBR pathway. There was no significant difference in ET-1 secretion between ETAR inhibition group and control group. **p<0.01 versus control. C. ET-1/ETBR pathway inhibitor or si-ETBR suppressed protein level of p-eNOS and increased protein level of p-p65. Therefore, eNOS might be related with NF-kappaB. D. si-eNOS significantly increased protein level of p-p65, indicating p-p65 was regulated by eNOS. **p<0.01 versus WT. E-F. Compared with si-eNOS group, mRNA expression and secretion of ET-1 were significantly suppressed in si-eNOS+Bay group. **p<0.01 versus WT or si-eNOS.

### Inhibition of NF-kappaB pathway ameliorated DN in ETBR^−/−^ mice in vivo

Four indices (serum creatinine, urinary albumin, serum ET-1 and kidney ET-1) were detected in WT, eNOS^−/−^, ETBR^−/−^, and ETBR^−/^+Bay mice groups. As shown in Figure 8A-D, the four indices were significantly increased in eNOS^−/−^ group than that of WT group, and the promotion effects were similar as ETBR^−/−^ mice, which suggested that eNOS pathway had important role in the treatment of DN, and eNOS pathway was suppressed in ETBR^−/−^ mice, therefore, ETBR^−/−^ mice had higher ET expression. Inhibition of NF-kappaB pathway in ETBR^−/−^ mice decreased the four indices, which indicated that NF-kappaB pathway played important role in endothelial cells. We observed that ECM-related proteins in eNOS^−/−^ mice were higher than WT mice (Figure 8E). After inhibition of NF-kappaB pathway in ETBR^−/−^ mice, ECM-related proteins were decreased, which suggested NF-kappaB pathway could regulate ET-1 expression to suppress ECM generation in MC (Figure 8E). Moreover, PAS staining showed that enlargement of glomeruli was observed in STZ-diabetic mice, and enhanced glomerulosclerosis was present in STZ-diabetic eNOS^−/−^ and ETBR^−/−^ mice.

**Figure 8.**
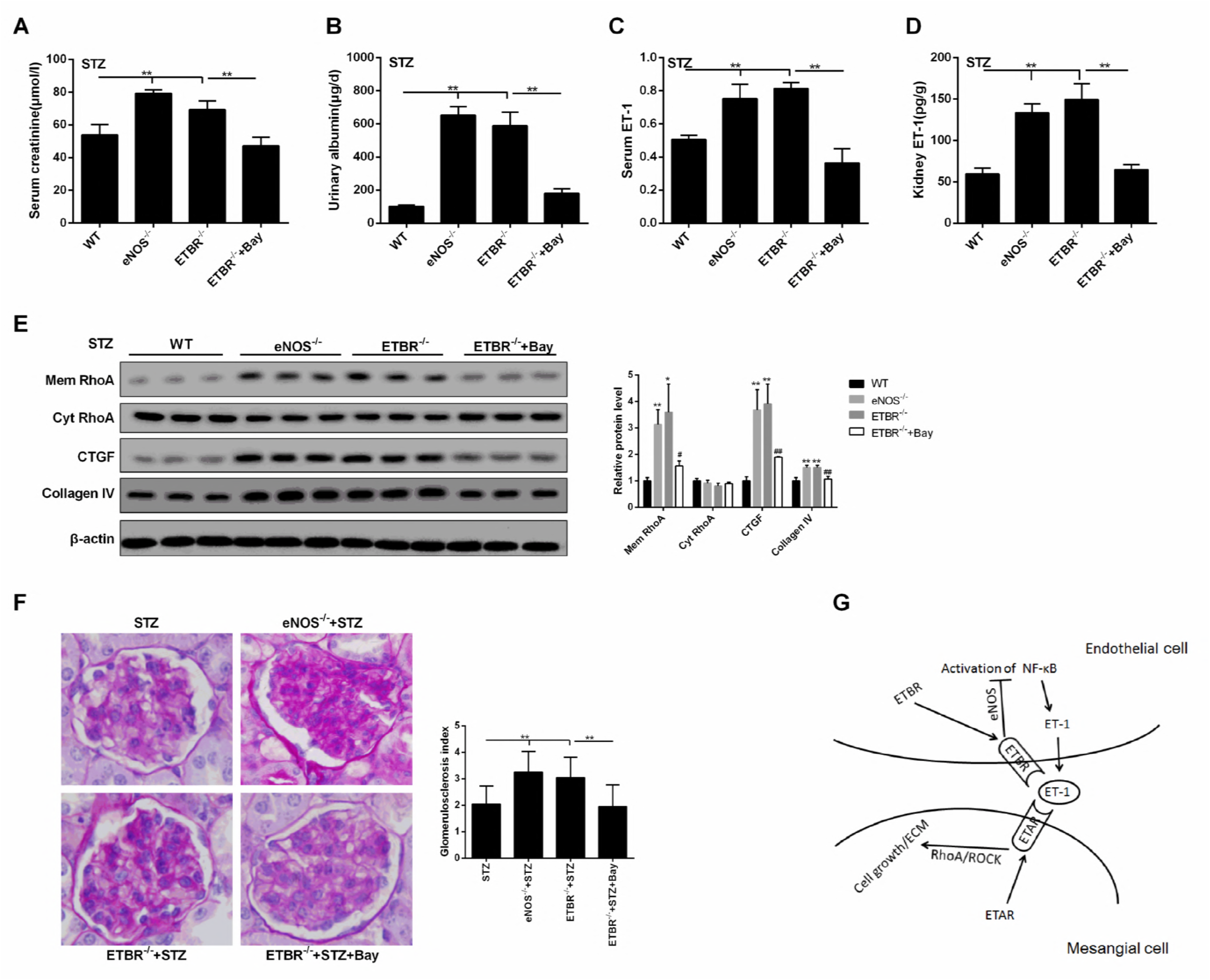
Inhibition of NF-kappaB pathway ameliorated DN in ETBR−/− mice in vivo. Bay was injected into mice between seventh and 10th weeks after STZ treatment. Mice were divided into four groups, namely WT, eNOS^−/−^, ETBR^−/−^, and ETBR^−/−^ Bay. N=3. A-D. Compared with WT mice, four indices (serum creatinine, urinary albumin, serum ET-1 and kidney ET-1) were increased in eNOS^−/−^ mice, and the promotion effects were similar as ETBR^−/−^ mice. This result suggested that eNOS pathway had important role in the treatment of diabetic nephropathy, and eNOS pathway was suppressed in ETBR^−/−^ mice, therefore, ETBR^−/−^ mice had higher ET expression. Inhibition of NF-kappaB pathway in ETBR^−/−^ mice decreased the four indice. **p<0.01 versus WT or ETBR^−/−^. E. ECM-related proteins (CTGF, collagen IV) in eNOS^−/−^ mice were higher than WT mice. After inhibition of NF-kappaB pathway in ETBR^−/−^ mice, ECM-related proteins were decreased, which suggested NF-kappaB pathway could regulate ET-1 expression to suppress ECM generation of MC. **p<0.01 versus WT or ETBR^−/−^. F. PAS staining showed that enlargement of glomeruli was observed in STZ-diabetic mice, and enhanced glomerulosclerosis was present in STZ-diabetic eNOS^−/−^ and ETBR^−/−^ mice. After inhibition of NF-kappaB pathway in ETBR^−/−^ mice, glomerulosclerosis was relieved. **p<0.01 versus STZ or ETBR^−/−^ +STZ. G. Cascade diagram of signaling pathways.

After inhibition of NF-kappaB pathway in ETBR^−/−^ mice, glomerulosclerosis was relieved (Figure 8F). These findings suggested that inhibition of NF-kappaB pathway ameliorated DN in ETBR^−/−^ mice, which indicated NF-kappaB pathway in endothelial cells could regulate ET-1. And eNOS^−/−^ mice had similar symptoms as ETBR^−/−^ mice, suggesting eNOS pathway in endothelial cells could regulate ET-1 production.

## Discussion

In the present study, we observed that STZ-diabetic ETBR^−/−^ mice had higher levels of renal damage signs (serum creatinine and urinary albumin), increased mRNA and protein levels of ET-1 and enhanced glomerulosclerosis *in vivo*. Besides, protein levels of CTGF, ETAR and p-p65 were upregulated, whereas protein level of p-eNOS was down-regulated in STZ-diabetic ETBR^−/−^ mice. Under high glucose condition, ETBR^−/−^ endothelial cells secreted large amount of ET-1, promoted MC proliferation, ECM accumulation of MC *in vitro*. We further demonstrated that ET-1 over-expression in ETBR^−/−^ endothelial cells was regulated by NF-kapapB pathway, and found ET-1/ETBR suppressed NF-kappaB via eNOS to modulate ET-1. We also proved ET-1/ETAR promoted RhoA/ROCK pathway in MC, thus to modulate MC proliferation and ECM accumulation. Based on the crosstalk between endothelial cells and MC, our results first revealed that in STZ-diabetic ETBR^−/−^ mice, ET-1 binding to ETBR was suppressed, so NF-kappaB pathway was activated via inhibiting eNOS, thus to secrete large amount of ET-1. Then, ET-1 binding to ETAR in MC, so RhoA/ROCK pathway was promoted, thus to accelerate MC proliferation and ECM accumulation.

Many studies have indicated that eNOS deficiency in diabetic mice aggravated renal injury and contributed to the pathogenesis of DN through blocking vascular endothelial growth factor receptor (VEGFR) to affect urinary albumin development, or blocking renin-angiotensin system (RAS) to control blood pressure[14, 15]. Yuen et al pointed out that under HG condition, eNOS deficiency can change the secretory reaction of endothelial cells [15]. Due to the activation of p-Akt, eNOS was activated in endothelial cells which led to phosphorylation of eNOS and NO release [16]. Liu et al found that ETBR could cause eNOS to be phosphorylated and induced NO production in sinusoidal endothelial cells via PI3K/Akt pathway [17]. Our results were consistent with the previous reports, showing that inhibition of ET-1/ETBR reduced p-eNOS expression.

NF-kappaB, a transcription factor that has five subunits, namely p50, p52 RelA/p65, c-Rel and RelB, plays a critical role in inflammatory process and metabolic disease [18]. It has been reported that high blood glucose, urinary albumin, angiotensin II could contribute to NF-kappaB activation [19, 20]. Evidence have shown that NF-kappaB activation in endothelial cells exerted a vital role in DN. Suppression of NF-kappaB attenuated HG-induced endothelial cell inflammation [21]. Advanced glycation end products increased NF-kappaB-binding activity to the promoter of ET-1 thus to increase ET-1 expression [22]. Researchers reported that C-peptide protected DN by preventing NF-kappaB from recruiting p300 and binding to the inos promoter [23]. Kolati et al demonstrated that NF-kappaB inhibitor BAY 11-7082 ameliorated DN by inhibiting renal inflammation and oxidative stress [24]. Moreover, glycolipids suppressed LPS-induced phosphorylation of NF-kappaB and promoted eNOS activation, and NOS inhibitor reduced glycolipids-induced NF-kappaB suppression [25], which suggested eNOS could negatively modulate NF-kappaB. However, the effect of eNOS/NF-kappaB pathway on ET-1 expression was not revealed in endothelial cells exposed to HG. Our results showed NF-kappaB regulated promoter of ET-1 at transcription level, and the expression of p65, a constituent of NF-kappaB, was remarkably upregulated after ET/ETBR inhibition or eNOS silence. Besides, si-eNOS remarkably increased ET-1 expression and Bay reversed the improvement of ET-1. This study first revealed suppression of ET-1/ETBR largely upregulated ET-1 expression through eNOS/NF-kappaB pathway in ETBR^−/−^ endothelial cells exposed to HG.

RhoA is a protein that cycle between active and inactive forms depending on binding to GTP or GDP [26]. ROCK, a serine/threonine kinase, is a downstream target of ROCK. Studies have shown that HG activated RhoA/ROCK pathway in MC, which contributed to the progression of DN [27, 28]. Researchers proved ET-1 binding to ETAR was involved in CTGF synthesis via RhoA/ROCK pathway in vascular smooth muscle cells [29]. Lee et al further discovered that over-expression of ET-1 binding to ETAR facilitated RhoA/ROCK pathway to induce collagen synthesis and proteinuria in hypertensive rats [30]. However, supporting evidences are still lacked for the acceleration effect of ET-1 binding to ETAR on MC proliferation and ECM accumulation via RhoA/ROCK pathway. Our result showed that membrane ETAR was highly expressed after ET-1 treatment, and inhibition of ET-1/ETAR suppressed MC proliferation, decreased CTGF and collagen IV protein levels, and inhibited RhoA/ROCK pathway. Moreover, RhoA/ROCK inhibitor suppressed MC proliferation, decreased ECM accumulation and promoted cell apoptosis with the treatment of ET-1. Therefore, we first proved that under HG condition, large amount of ET-1 binding to ETAR accelerated MC proliferation and ECM accumulation through promoting RhoA/ROCK pathway.

In conclusion, we have demonstrated that in HG exposed ETBR^−/−^ endothelial cells, suppression of ET-1 binding to ETBR activated NF-kappaB pathway via inhibiting eNOS, thus to secrete large amount of ET-1. Due to the crosstalk between endothelial cells and MC, ET-1 binding to ETAR in MC promoted RhoA/ROCK pathway, thus to accelerate MC proliferation and ECM accumulation. Therefore, STZ-diabetic ETBR^−/−^ mice accelerated the progression of DN.

